## Supplemental Figures for "Reciprocal regulation between cell mechanics and ZO-1 guides tight junction assembly and epithelial morphogenesis"

Alexis J. Haas *et al.*

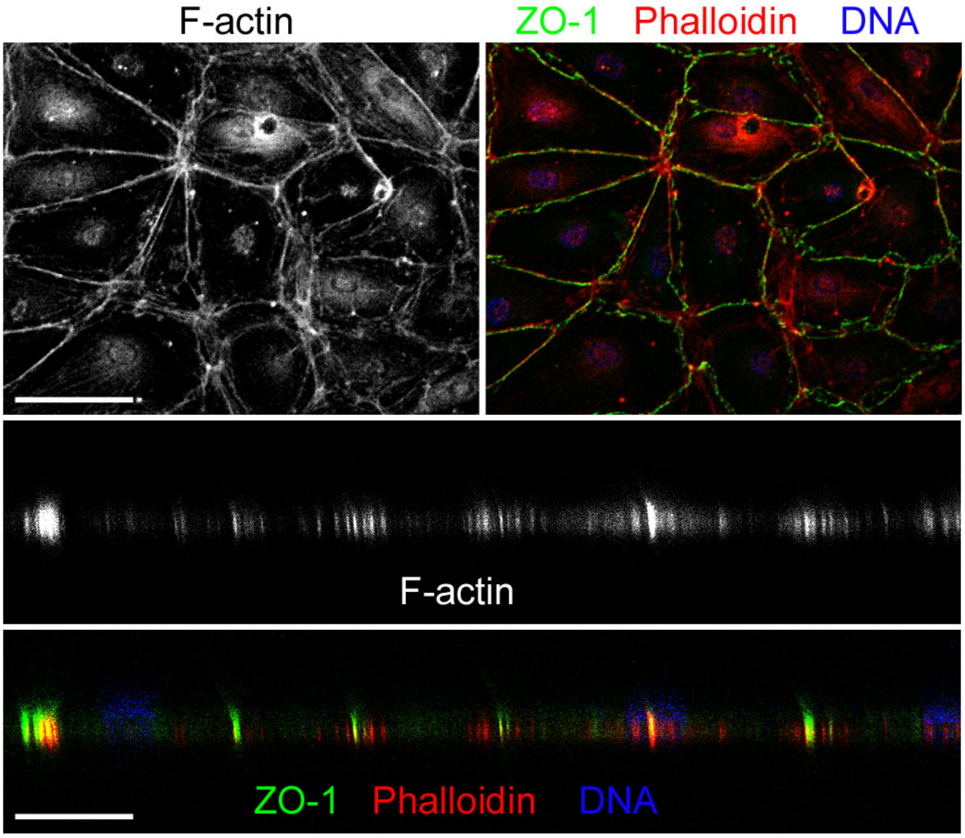


**Figure S1. F-actin distribution in primary microvascular endothelial cells.**
Endothelial cells were seeded on Matrigel-coated glass and immunostained for ZO-1. F-actin was labelled with fluorescent phalloidin and nuclei with Hoechst dye. Cells were visualized in xy (top panel) and in xz orthogonal cross-sections (middle and bottom panels). Magnification bars, 20 μm.

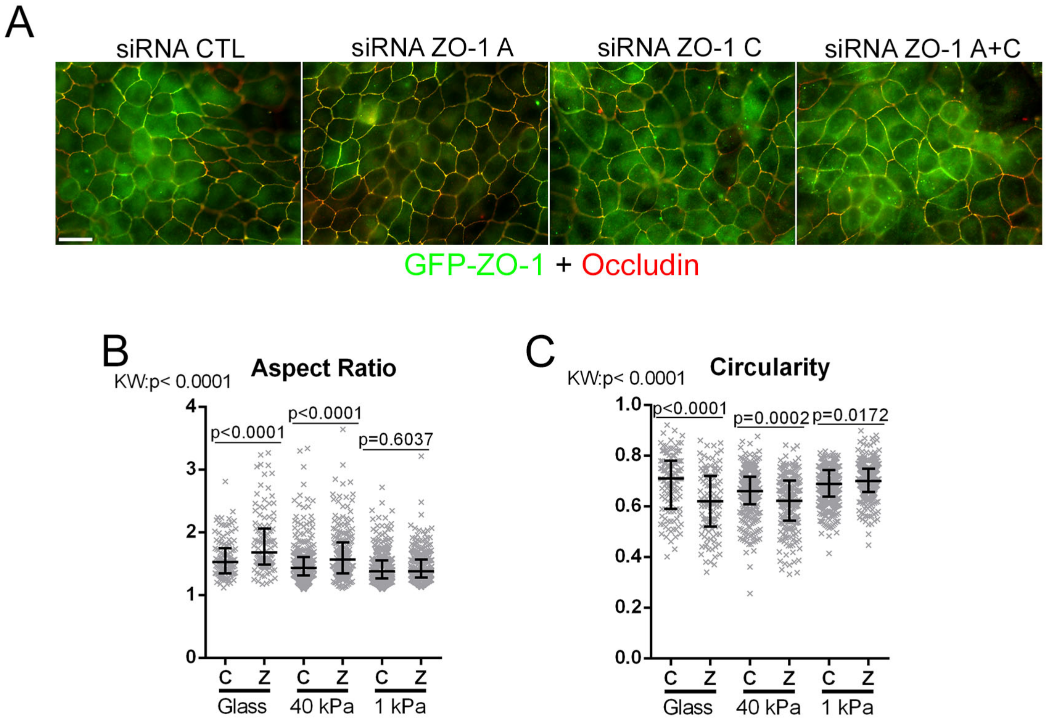


**Figure S2. Expression of GFP-mZO-1 and soft ECM rescue the ZO-1 depletion phenotype.**
**A** GFP-mZO-1 expressing MDCK cells were transfected with siRNAs as indicated and then plated on glass followed by immunostaining for occludin. Images show overlays of GFP and occludin images. **B, C** Wild-type MDCK cells transfected with control (c) or pooled siRNAs against ZO-1 (z) were seeded on glass or hydrogels (40 or 1kPa). Junctional immunostaining was then performed, and the resulting images were used to segment individual cells and perform a morphometric analysis of aspect ratios (B) and circularity (C). Data points represent analysed cells. Magnification bar, 20 μm.

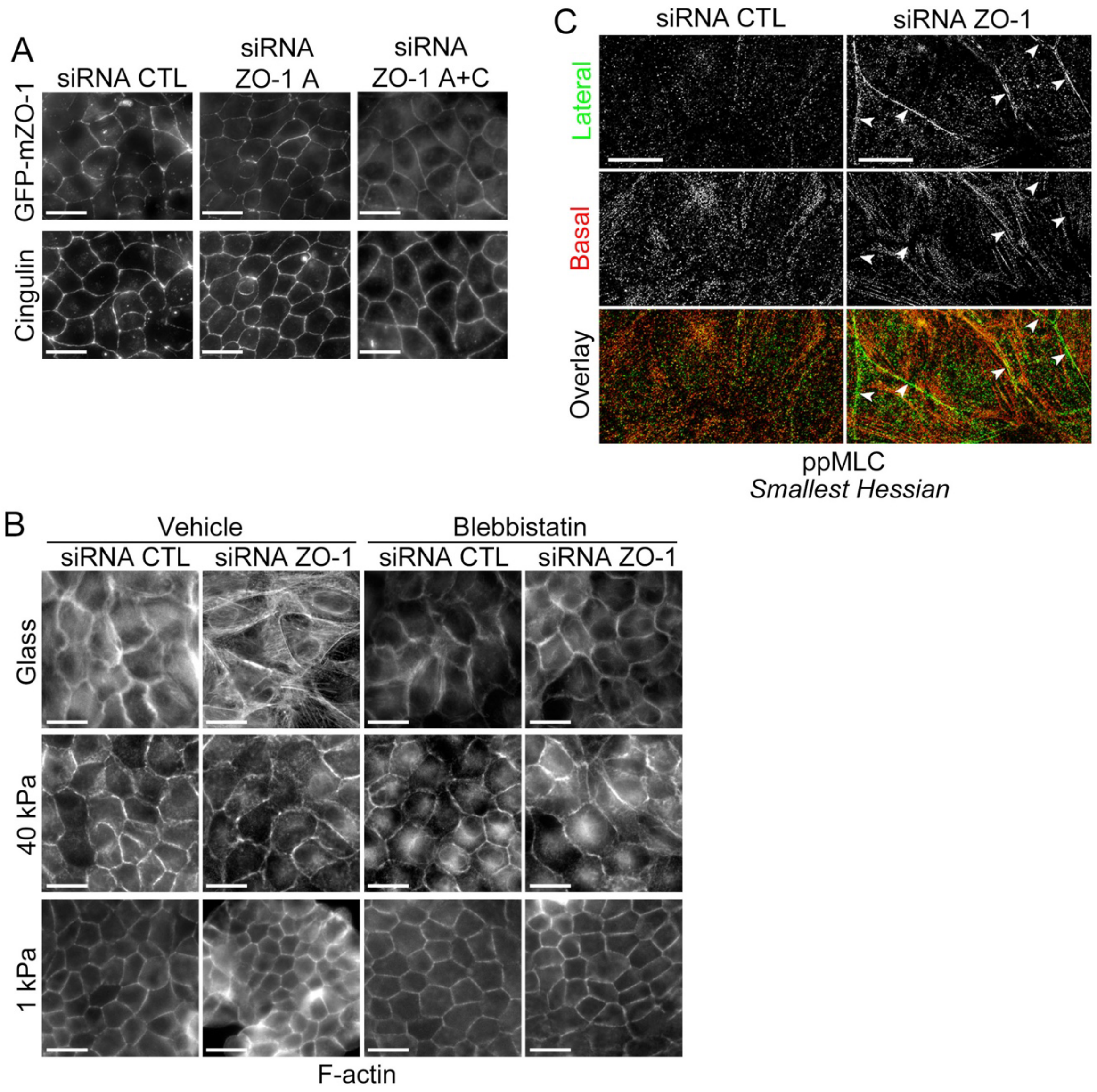


**Figure S3. Junction formation rescue and actomyosin remodelling upon ZO-1 depletion.**
**A** The GFP-mZO-1 cell line was transfected with siRNAs as indicated before immunolabelling cingulin and imaging. **B, C** siRNA transfected MDCK cells were seeded on glass, or 40 kPa or 1 kPa hydrogels. After cell fixation, cells were labelled using phalloidin to visualize F-actin (B) or ppMLC (C, cells gown on glass). In panel C, microscopy images were processed using the smallest Hessian differentiation operation to enhance the visual discrimination between basal and lateral features of actomyosin. Magnification bars, 20 μm.

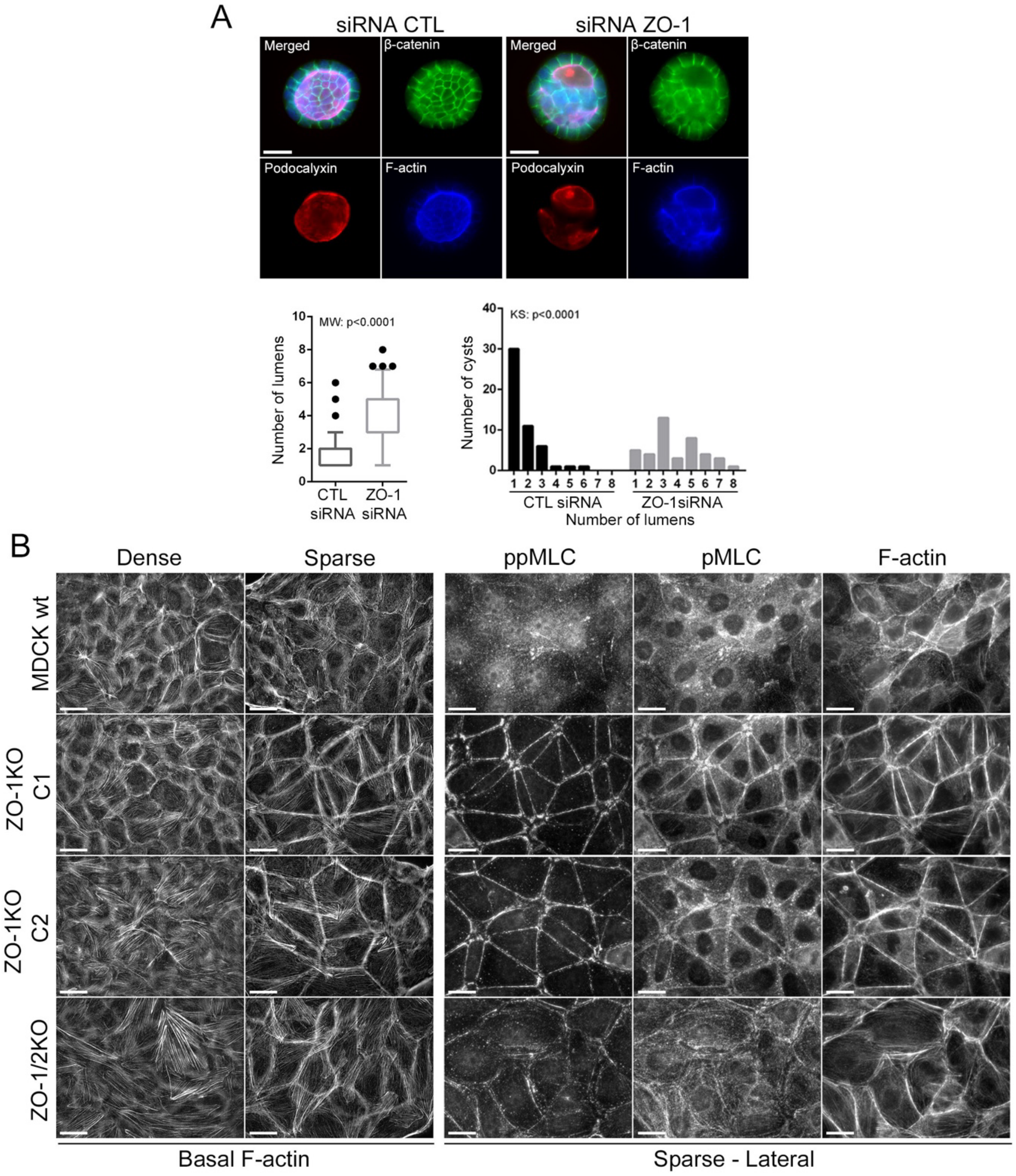


**Figure S4. ZO-1 deficiency disrupts 3D morphogenesis and stimulates actomyosin remodelling.**

**A** siRNA transfected MDCK cells were seeded in a matrix consisting of Matrigel and collagen I to form 3D spheroids. After fixation the cysts were stained as indicated, the number of lumens per cyst was manually counted, and the distribution of the cysts population was plotted over the number of lumens per cyst counted in each condition. **B** Wild-type and knockout MDCK cells were seeded on glass or hydrogels (40 kPa and 1 kPa) before fluorescently staining F-actin, pMLC and ppMLC. The most in-focus basal slices from acquired z-stacks were enhanced to visualize stress fibres in sparse and dense areas with the F-actin staining (left panels). Images showing lateral focal planes of sparse areas are shown for pMLC, ppMLC and F-actin (right panels). Magnification bars, 20 µm.

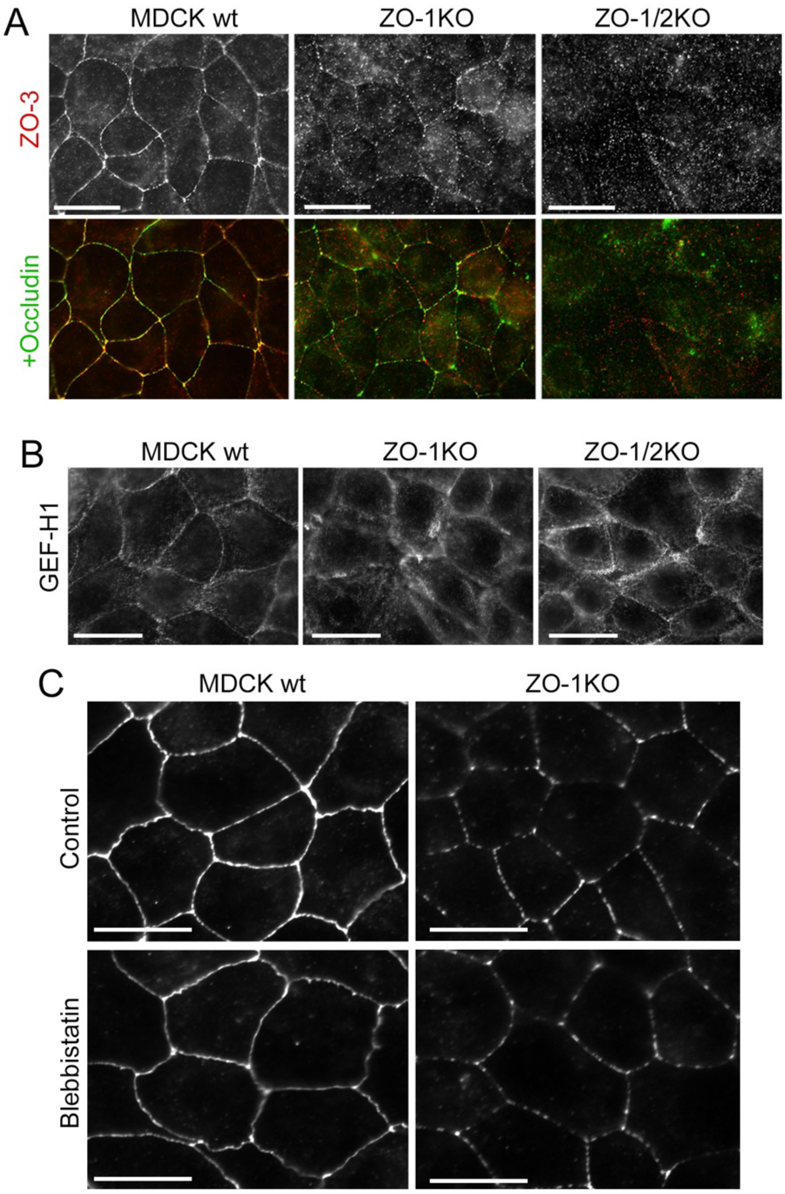


**Figure S5. TJ assembly analysed in filter-grown cells.**
Wild-type and knockout MDCK cells labelled with antibodies against the indicated proteins were imaged by bright field microscopy.

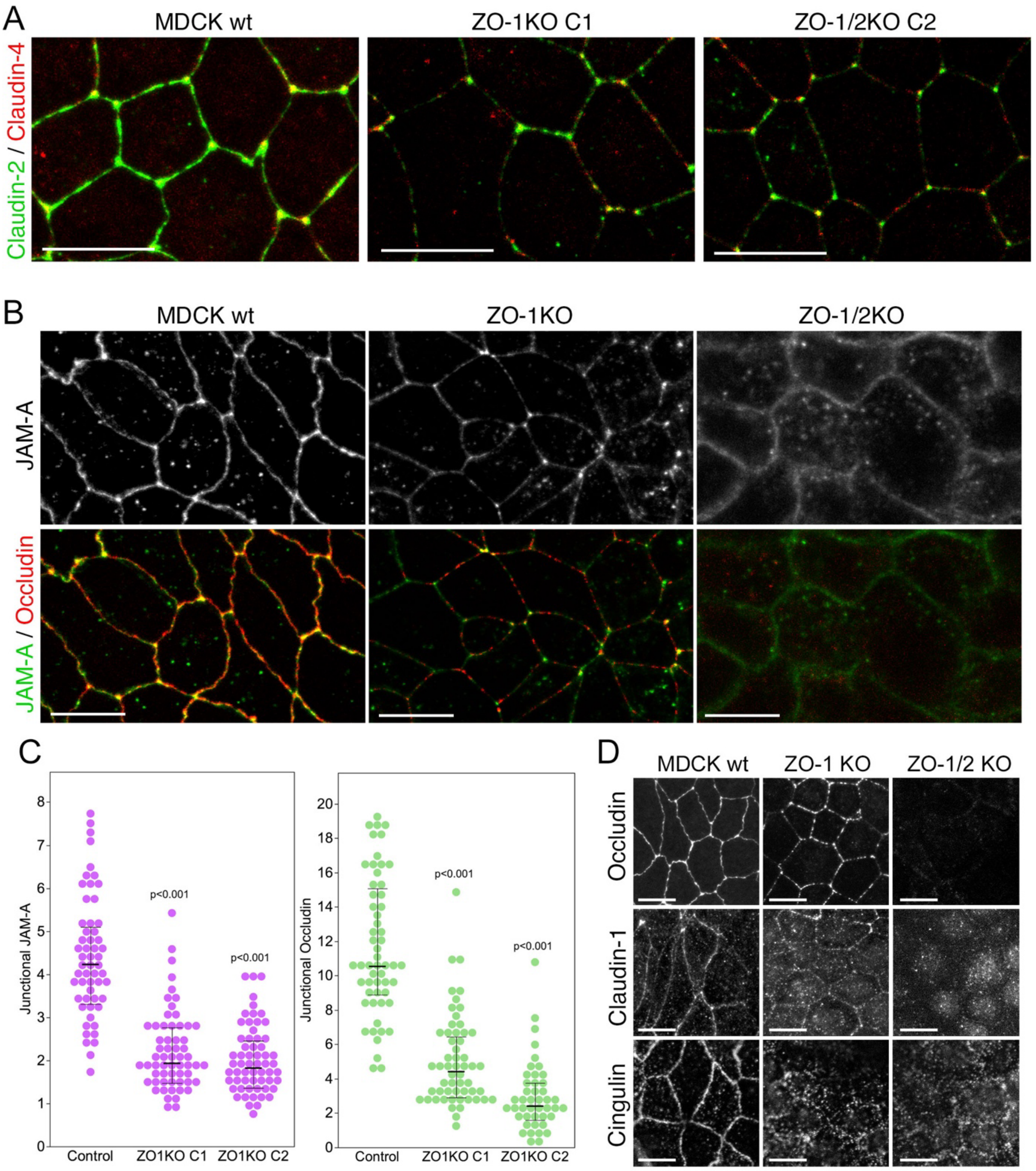


**Figure S6. TJ assembly analysed in filter-grown cells.**
Wild-type and knockout MDCK cells were grown on filters prior to analysis by immunofluorescence and confocal microscopy. All confocal XY sections are maximum intensity projections of all sections containing junctional staining. The fluorescence intensities of JAM-A and occludin at cell junctions were quantified in z-sections (C). Magnification bars, 20 µm.
